# Integrated Transcriptome and Metabolome Analyses Reveal the Mechanism Regulating Bulbil Initiation and Development in *Cystopteris chinensis*

**DOI:** 10.1101/2024.09.18.613657

**Authors:** An Yu, Xiaohong Chen, Wenkai Xi, Xia Zhao, Yazhu Wang, Zhihong Gong, Xiaofeng Zhou

**Affiliations:** College of Forestry, Sichuan Agricultural University; Key Laboratory of Ecological Forestry Engineering of Sichuan Province, College of Forestry, Sichuan Agricultural University; Bijie Forestry Bureau; Qingchuan Forestry Bureau

**Keywords:** *Cystopteris chinensis*, fern, bulbil, transcriptome, metabolome, development

## Abstract

*Cystopteris chinensis* is an endangered fern endemic to China, which only has a small wild population due to its poor reproductive ability. However, we recently found that it can produce bulbils on its pinnule to generate new plants but the molecular mechanism underlying this unique phenomenon remained unknown. In this study, we have identified four distinct stages in the initiation and development of bulbils based on morphological and anatomical observation. We performed transcriptome and metabolome analyses on the collected samples at each stage. Through KEGG enrichment analysis, it was found that the phytohormone signal transduction, starch and sucrose metabolism, phenylpropanoid biosynthesis, and flavonoid biosynthesis pathways play a significant role in regulating bulbil initiation and development. Specifically, the involvement of three phytohormones and sugar substances was identified in the process of bulbil initiation. Our study provides the first detailed observation of the bulbils in *C. chinensis* and explains their initiation and development at the molecular level. However, more in-depth studies are needed to discover the functions of key genes controlling the formation of bulbils to conserve the endangered *C. chinensis* population.

## Introduction

Free-sporing vascular plants are classified into two distinct evolutionary lineages: lycophytes and ferns (Kenrick and Crane, 1997). Ferns, conversely, are a more diverse and the second largest group of vascular plants after the flowering plants (Ranker and Haufler, 2008). The swift proliferation of ferns has been extensively recorded in the geological archives, particularly following significant extinction episodes (Thomas and Cleal, 2022). Before angiosperms took over terrestrial ecosystems, ferns, bryophytes, lycophytes, and gymnosperms dominated with over 10,000 species in a range of shapes and sizes (Schneider *et al*., 2004). Phylogenetic studies have revealed that horsetails and ferns are the most closely related plant species to seed plants (Pryer *et al*., 2001). Being an important part of vascular higher plants, the knowledge of fern biology is the key to understanding the evolution of land plants (Li *et al*., 2018). Ferns and lycophytes are unusual plants with a gametophyte and sporophyte life cycle, with the sporophyte usually being the dominant stage (Haufler *et al*., 2016). Other than reproducing through spores which is sexual propagation, ferns can propagate rapidly using tissue (leaf, stem, root, bulbil) cultured in vitro (Higuchi *et al*., 1987; Higuchi and Amaki, 1989).

A bulbil, also known as a bulblet, is distinguished by the presence of a petite bud attached to a brief stem (D. Bell and Bryan, 2008). This unique structure allows the bulbil to develop roots and eventually grow into a new plant independently from the parent plant (Arizaga and Ezcurra, 2002; Wang and Cronk, 2003). Research on the formation of bulbils has been limited to a few plant species, including *Dioscorea polystachya*, *Dioscorea batatas*, *Allium sativum*, *Titanotrichum oldhamii*, *Lilium sulphureum*, *Pinellia ternata*, *Lilium lancifolium, Kalanchoë* and *Agave tequilana* (Zhang *et al*., 1994; Wang and Cronk, 2003; Garcês *et al*., 2007; Abraham-Juárez *et al*., 2010; Walck *et al*., 2010; Li *et al*., 2012; Luo *et al*., 2014; Yang *et al*., 2017). In *L. lancifolium*, the formation of bulbils may be governed by genetic factors associated with hormone signaling pathways and starch biosynthesis (Yang *et al*., 2017). Similarly, research on *L. davidii* var. *unicolor* indicates that sucrose plays a critical role in bulbil morphogenesis, whereas starch is essential for bulbil emergence and growth (Li *et al*., 2014). In *T. oldhamii* from the Gesneriaceae family, *FLORICAULA* (*GFLO*) has been identified as a potential regulatory gene involved in bulbil formation (Wang *et al*., 2004). Moreover, the expression of two class I *KNOX* genes in *A. tequilana* is observed during bulbil initiation and increases as the bulbils reach maturity (Abraham-Juárez *et al*., 2010). Despite the identification of numerous genes involved in bulbil initiation and development, the metabolic pathways and complex regulatory mechanisms of bulbil production remain unknown.

*Cystopteris chinensis* (Ching) X.C. Zhang & R. Wei, an endangered fern species of the Cystopteridaceae family, is an endemic fern to China (R.C., 1966; Wei and Zhang, 2014). Its habitat only appears to be a temperate broadleaf forest of Erlang Mountain and Emei Mountain in Sichuan, China (Zhang, 2012). In China, all fern species of *Cystopteris* exhibit notable genetic differences from *C. chinensis*, whereas *Cystopteris bulbifera* (L.) Bernh. from the eastern United States is closely related as a sister group to *C. chinensis* (Wei and Zhang, 2014). The severe decline in the population of *C. chinensis* can primarily be attributed to habitat destruction and reduced spore reproductive capabilities (Cai *et al*., 2016). Research on artificial in vitro tissue culture of the spores revealed the species’ strict substrate needs. Germination is fastest at 20°C on its native soil. Even then, the highest germination rate is 57.8% (Cai *et al*., 2016; Chen *et al*., 2016).

Our recent discovery regarding the reproductive capabilities of *C. chinensis*, specifically its ability to form bulbils at the apex of its pinnules and efficient propagation through tissue culture techniques, has provided valuable insights. Four distinct morphological stages in the development of bulbils can be observed. In this study, we delve into the molecular aspects of this process by analyzing transcriptome profiles of bulbils at different differentiation stages and monitoring changes in metabolites during bulbil development, we attempt to understand the molecular mechanisms involved in bulbil initiation and development of *C. chinensis*, and to set the stage for future investigations into its reproductive biology and conservation strategies.

## Materials and methods

### Plant Materials

We used the wild population of Tuan niuping in Erlang Mountain, Sichuan Province, China, the native type locality of the *Cystopteris chinensis*, as the experimental material. Tissue samples were collected regularly from January 2022 to January 2023, and subjected to histological examinations using a stereo microscope and transmission electron microscope to observe and set the four key developmental stages of bulbil (**Figure 1A-D**), involving S1 (absence of bulbils), S2 (initial appearance), S3 (bulbils differentiation), and S4 (full maturation). From April to May 2023, we sampled field populations at average intervals of 15 days, three biological replicates for 12 samples were produced from each stage to validate the research conclusions. The samples were immediately frozen in liquid nitrogen at -80°C for further transcriptome and metabolome analyses utilizing RNA-Seq, ultra-high performance liquid chromatography-mass spectrometry (UHPLC-MS).

**Fig. 1.**
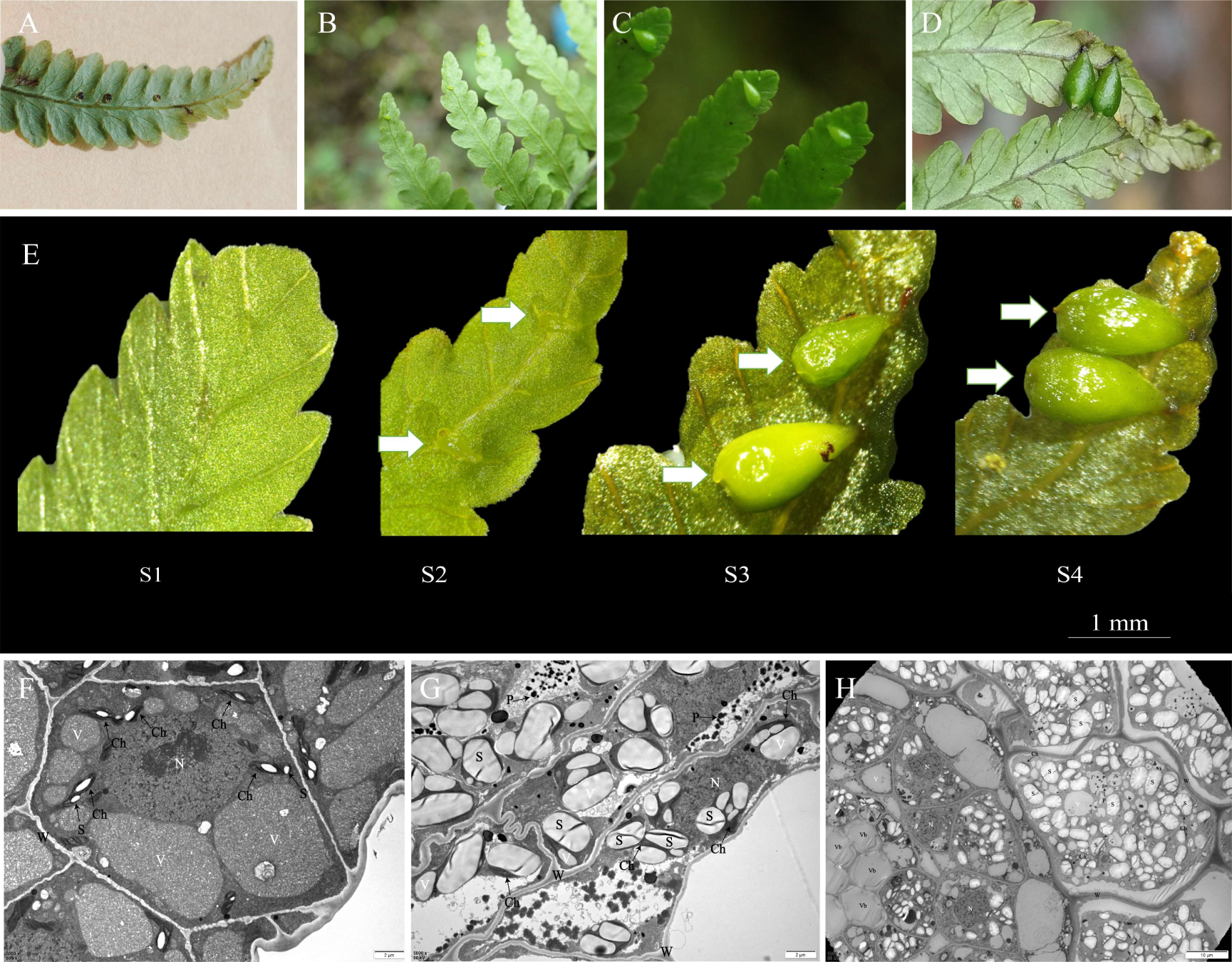
(A, E) Pinnule without bulbil. (B, E) Initiation of bulbil. (C, E) Differentiation of bulbil. (D, E) Maturation of bulbil. (F) Ultrastructure of bulbil in S2. (G) Ultrastructure of bulbil in S3. (H) Ultrastructure of bulbil in S4. Ch: chloroplast; P: pigment granules; S: starch; N: nucleus; V: vesicles; Vb: vascular bundle; W: cell wall.

### Ultrastructural observation

The bulbils at various stages of growth were carefully examined using transmission electron microscopy (TEM). Prefixed with a 3% glutaraldehyde, then the tissue was postfixed in 1% osmium tetroxide, dehydrated in series acetone, infiltrated in Epox812 and embedded. The semithin sections were stained with methylene blue and the Ultrathin sections were cut with a diamond knife, and stained with uranyl acetate and lead citrate. Samples were prepared following Trizro’s method (Tizro *et al*., 2019), and image acquisition was performed with a JEM-1400FLASH transmission electron microscope by Nippon Electronics (Engel and Colliex, 1993).

### RNA extraction and RNA-Seq

Total RNA was extracted from four different stages on a cBot Cluster Generation System using TruSeq PE Cluster Kit v3-cBot-HS (Illumia) according to the manufacturer’s instructions. To select cDNA fragments of preferentially 370∼420 bp in length, the library fragments were purified with the AMPure XP system (Beckman Coulter, Beverly, USA). RNA integrity and the library quality were assessed on the Qubit2.0 Fluorometer by using the RNA Nano 6000 Assay Kit of the Bioanalyzer 2100 system (Agilent Technologies, CA, USA). Total 24 cDNA libraries, including S1_1, S1_2, S1_3, S2_1, S2_2, S2_3, S3_1, S3_2, S3_3, S4_1, S4_2 and S4_3, were constructed and sequenced using Illumina HiSeq™ 4,000 platform at Novogene Bioinformatics Technology Co. Ltd. (Beijing, China).

### Data filtering, transcript assembly, and gene functional annotation

We prepared clean reads from the sequencing data by eliminating reads with adapters, those with over 10% unknown nucleotides and low-quality reads were deleted. We then assessed the Q20, Q30, GC content, and sequence duplication levels using the clean data. Lastly, we carried out a de novo assembly based on the clean data using the Trinity program (Grabherr *et al*., 2011). To gain a comprehensive understanding of the functional information, we aligned all genes using BLAST against several databases with an e-value threshold of 1e^-05^. The protein sequences of unigenes with the highest similarity were retrieved for functional annotation and classification.

### Analysis of Differential Gene Expression

In our research, we divided all samples into 12 comparisons (S2vs.S1, S3vs.S1, S4vs.S1, S3vs.S2, S4vs.S2, S4vs.S3, S2vs.S1, S3vs.S1, S4vs.S1, S3vs.S2, S4vs.S2, S4vs.S3). This was done to thoroughly identify the significant genes related to bulbil initiation and development. We determined correlations by calculating Pearson’s correlation coefficient. Then we calculated the expression of unigenes using the FPKM (fragments per kilobase per million base pairs) method. The DEseq2 package (1.20.0) in R was used to analyze unstandardized read count data between two samples, using a false discovery rate (FDR) < 0.05, Benjamini and Hochberg’s approach, and absolute log2 FC≥1 to identify differentially expressed genes. The cluster Profiler R program was used to analyze GO enrichment of DEGs using Wallenius non-central hypergeometric distribution (corrected p<0.05) (Young *et al*., 2010). We performed KEGG (http://www.genome.jp/kegg/) enrichment analyses (p < 0.05 and FDR < 0.05) using cluster Profiler R package to test the statistical enrichment of differential expression genes (Wu *et al*., 2021).

### Metabolites Extraction and UHPLC-MS/MS Analysis

Each bulbil stage’s 100 mg of tissue was homogenized with liquid nitrogen and resuspended in prechilled 80% methanol by vortexing. After 5 minutes on ice, the samples were centrifuged at 15,000 g, 4°C for 20 minutes. The supernatant was diluted with LC-MS grade water to 53% methanol. The samples were then centrifuged at 15,000 g, 4°C for 20 minutes in a new Eppendorf tube. Finally, the supernatant was injected into the LC-MS/MS system for analysis (Want *et al*., 2013). Novogene Co., Ltd. (Beijing, China) performed UHPLC-MS/MS analyses utilizing a Vanquish UHPLC system (ThermoFisher, Germany) and an Orbitrap Q ExactiveTM HF or HF-X mass spectrometer. The samples were injected into a 100x2.1 mm Hypersil Gold column (1.9 μm) with a 12 minutes gradient and 0.2 mL/min flow rate. In positive polarity mode, eluent A was 0.1% formic acid in water and B was methanol. Eluent A was 5 mM ammonium acetate at pH 9.0 while B was methanol for negative polarity mode. The gradient profile was 2% B for 1.5 min, 2-85% B for 3 min, 85-100% B for 10 min, 100-2% B for 10.1 min, and 2% B for 12 min. Q ExactiveTM HF mass spectrometer operated at 3.5 kV spray voltage, 320°C capillary temperature, 35 psi sheath gas flow rate, 10 L/min auxiliary gas flow rate, 60 S-lens RF level, and 350°C auxiliary gas heater temperature. (Schrimpe-Rutledge *et al*., 2016).

### Analysis of Metabolomic Data

From UHPLC-MS/MS raw data files, Compound Discoverer 3.3 CD3.3 aligned, selected, and quantified metabolites. The key parameters were minimum intensity, 5ppm real mass tolerance, and 30% signal intensity tolerance. Following peak intensity normalization to overall spectral intensity. Normalized data projected molecular formulas and matched peaks with mzCloud, mzVault, and MassList databases for qualitative and relative quantitative results. Statisticians utilized R (3.4.3), Python (2.7.6), and CentOS (6.6). Chemicals with a relative peak area exceeding 30% in QC samples were eliminated and non-normally distributed data was adjusted. Metabolite identification and quantification concluded. We discovered metabolites using KEGG, HMDB, and LIPID Maps. MetaX was used for PCA and PLS-DA (Wen *et al*., 2017). Statistical analysis involved t-tests for metabolites with VIP > 1, p-value < 0.05, and fold change ≥2 or ≤0.5. Volcano shows metabolites by log2(FoldChange) and -log10(p-value) using R’s ggplot2. Pheatmap clustering heatmaps utilizing metabolite intensities’ z-scores. Metabolite correlations were assessed using R (Pearson technique) and significance was determined using cor.mtest() (P < 0.05). Functional analysis of metabolites and pathways with KEGG. Metabolic pathway enrichment analysis used x/n > y/N ratios and p-value < 0.05 for statistical significance.

### Integrated Correlation Analysis of Transcriptomic and Metabolomic

We conducted a correlation analysis between differentially expressed genes and metabolites using the Pearson method to compute correlation coefficients (r^2^) and p-values. The top 5 metabolites were selected based on metabolomics enrichment results, and the top 10 genes were identified from transcriptomics data. A metabolite-transcript correlation network was constructed using the R mixOmics package. KEGG pathway enrichment analysis was performed for both metabolites and transcripts, focusing on pathways enriched by both types of molecules in the same comparisons. A scatter plot visualizing metabolite-transcript KEGG enrichment was generated using the R ggplot2 package. Finally, pathways enriched by common differentially expressed metabolites and genes were mapped onto the iPath (https://pathways.embl.de/) (Letunic *et al*., 2008).

## Results

### Morphological and Ultrastructural Observation

Changes in the overall appearance of the bulbils were observed under the stereo microscope (**Fig. 1E**). Since early May 2023, *C. chinensis* pinnules were at stage S1, with smooth backs lacking spores or bulbils. Stage S2 bulbils of *C. chinensis* developed light-green, virtually translucent, elongated, teardrop-shaped bulbils at the middle of the pinnule’s apical lateral veins in mid-May. The bulbils’ parenchyma cells were tightly arranged, without cell gaps, and small, square or polyhedral. The nucleus was large, located in the center of the cell, and the nucleolus was also large. The cytoplasm was thick, and the vesicles were small and dispersed. The cytoplasmic organelles were rare, and a few chloroplasts were visible (**Fig. 1F**). In early June, *C. chinensis* entered stage S3, where bulbils grew, turned green, hardened, and produced spores. The bulbil parenchyma cells were the same as stage S2, but they were larger and had more chloroplasts, which were large and had starch granules, mitochondria, rough endoplasmic reticulum, and many pigment granules (**Fig. 1G**). At the end of June, it entered the maturity stage S4, and the form of bulbils reached the maximum, the color became dark green, the tip appeared to be concave and there were two small protrusions, The number of starch grains peaked and vascular bundles appeared in the middle of the cell (**Fig. 1H**) (**Supplementary Fig S12**). Histologically, the cells were loosely arranged, the cell volume increased, the plasma wall was separated, the cell membrane and cell wall ruptured, chloroplast morphology and structure in the cytoplasm disintegrated, and the starch granules grew larger (**Supplementary Fig S11**), and more pigment granules were seen in the cytoplasm. In conclusion, during the development of the bulbils, the cell volume and the volume of starch granules in the chloroplasts gradually increase.

### Dynamic transcriptomic profiles

No reference genome sequences for *C. chinensis* were available, thus transcripts were assembled de novo. To discover candidate genes for *C. chinensis* bulbil initiation and development, 12 libraries were created at four stages. Each library has almost 6.5 Gb of data and 277,836,575 raw reads. All sequencing data had Q20 and Q30 over 97% and 93%, respectively, with base mismatch rates of 0.03 (**Supplementary Table S1**). The data further indicated that the duplicate samples displayed a robust correlation (**Supplementary Fig. S1**). After filtering the raw data, sequencing error rate, and GC content distribution, 272,702,192 clean reads were collected for analysis. DEGs were identified based on |log2(Fold Change)| ≥ 1 and p-value < 0.05 (**Fig. 2A**). A total of 7,837 unigenes were obtained from these databases, which shows fewer genes in the S1 (1113 unigenes) and S2 (1516 unigenes) than in the S3 (1270 unigenes) and S4 (3938 unigenes) libraries (**Fig. 2B**). S2vs.S1, S3vs.S1, S4vs.S1 had six shared DEGs (**Fig. 2C**). While S3vs.S2 and S4vs.S2 had 2195 shared differential genes (**Fig. 2D**).

**Fig. 2.**
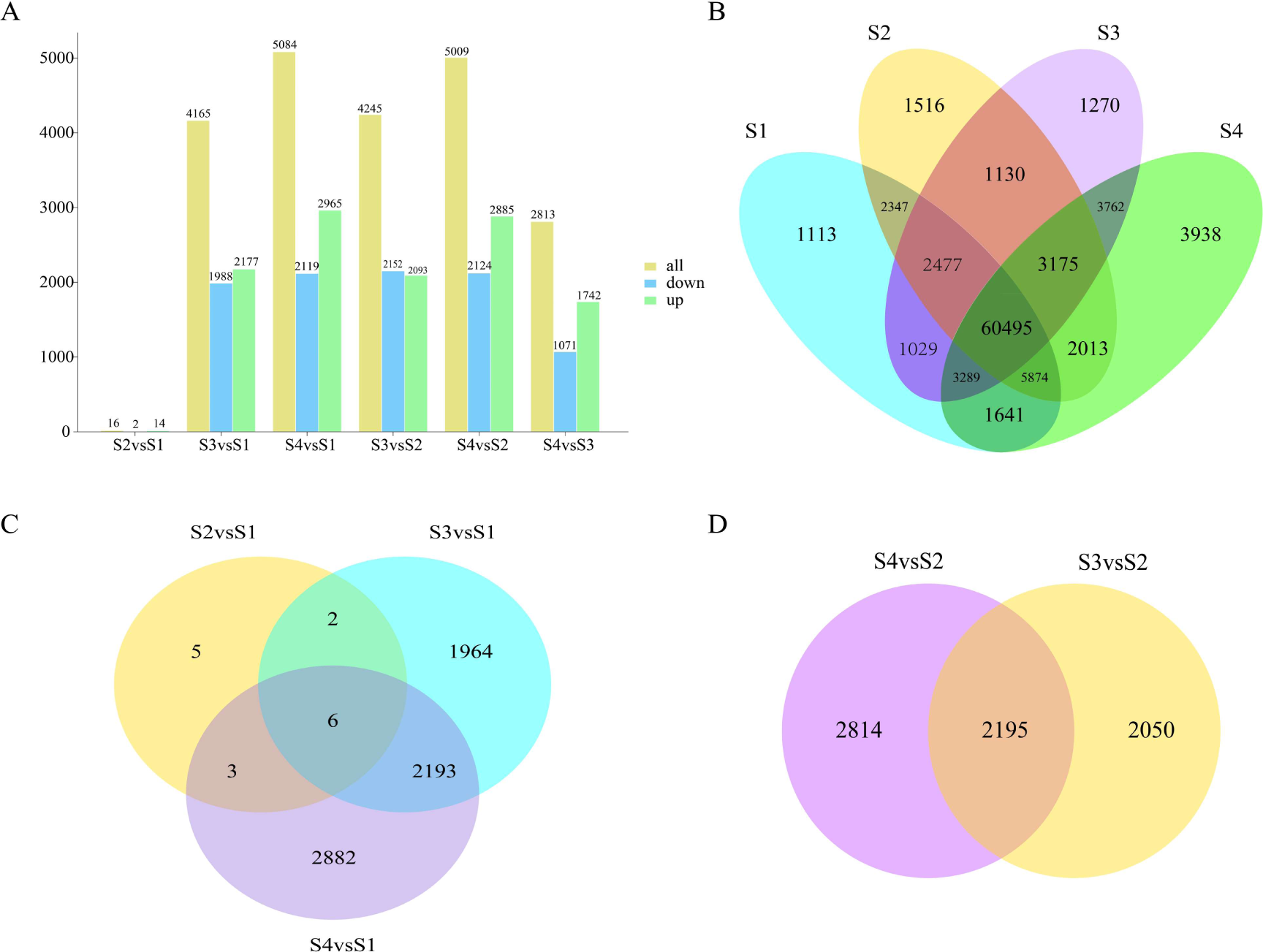
(A) Comparison of the number of up and down-regulated genes between different stages. (B) Venn diagram illustrating the unique and shared expressed genes in bulbil among different stages. (C) Venn diagram of DEGs among the three comparisons of bulbil initiation. (D) Venn diagram of DEGs among the two comparisons of bulbil development.

### Functional annotation of DEGs

DEG functions were classified by GO enrichment analysis. Padj < 0.05 indicated significant enrichment. Compared to samples from four stages, 177, 3534, 3785, 3561, 3699, and 3075 GO terms were enriched in six comparison groups(**Fig. 3, Supplementary Fig. S2**). It was found that in the comparison of the stages with and without bulbil (S2vs.S1, S3vs.S1, S4vs.S1), the differences mainly existed in the “biological process” and they are primarily enriched in lipid metabolic process, carbohydrate metabolic process, and cellular component biogenesis, respectively, and S3 was significantly enriched in carbohydrate metabolic process in comparison with S1 (**Fig. 3A**). In addition, it was found that the differences between S4vs.S2 were in the “biological process” of carbohydrate metabolism, the “molecular function” of oxidoreductase activity, and the “biological process” of carbohydrate metabolism (**Fig. 3B**).

**Fig. 3.**
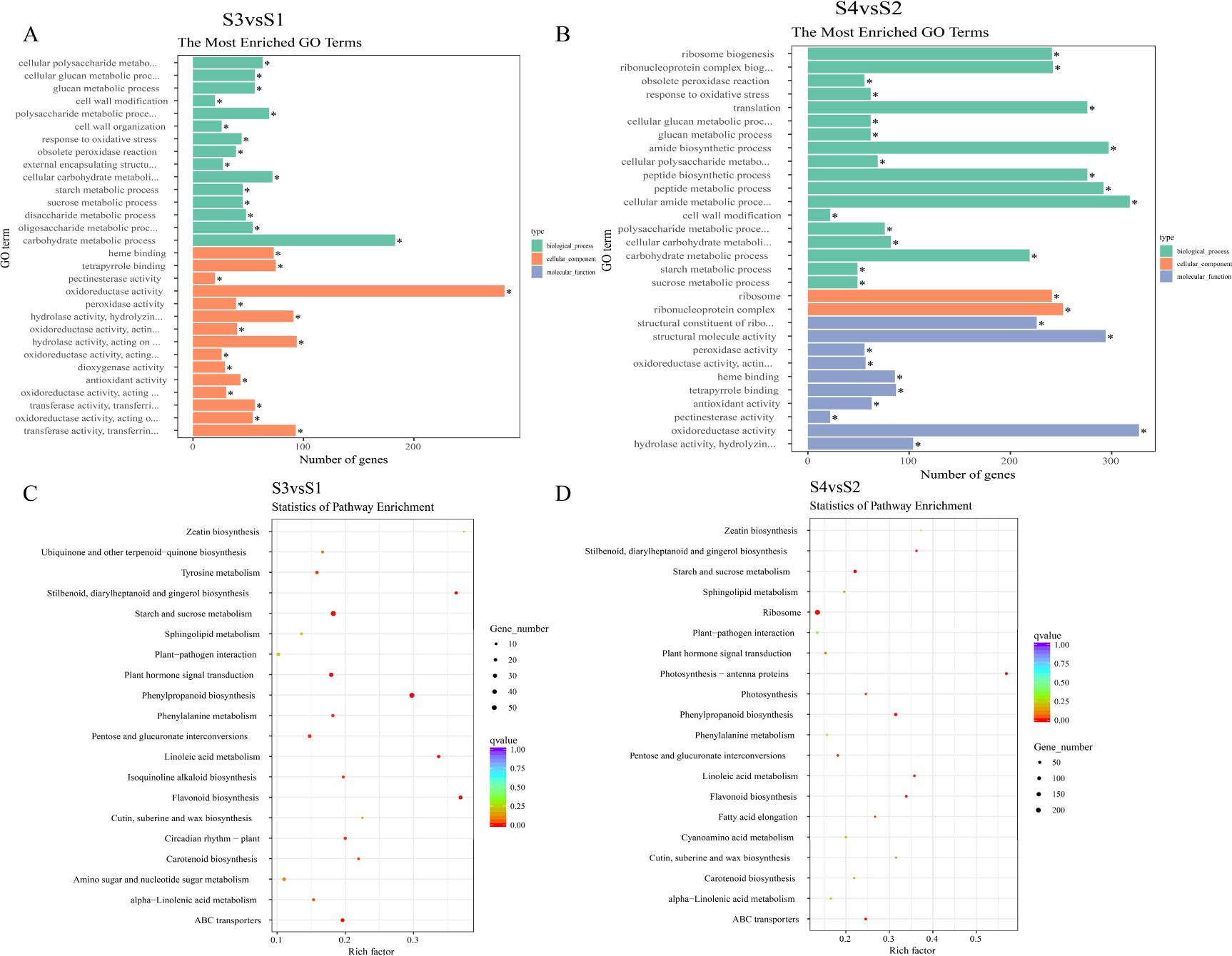
(A) GO annotation classification between S3vs.S1. (B) GO annotation classification between S4vs.S2. (C) Top 20 significantly enriched KEGG pathways related to initial bulbil in S3vs.S1. (D) Top 20 significantly enriched KEGG pathways related to bulbil development in S4vs.S2.

To investigate metabolic pathways, all DEGs were mapped to KEGG pathway enrichment analysis. Overall, 16 DEGs (S2vs.S1) were allocated to 4 pathways, 4165 DEGs (S3vs.S1) to 102 pathways, 5084 DEGs (S4vs.S1) to 99 pathways, 4245 DEGs (S3vs.S2) to 105 pathways, 5009 DEGs (S4vs.S2) to 101 pathways, and 2813 DEGs (S4vs.S3) to 93 pathways (**Supplementary Table S2**). A q<0.05 was used as the threshold to determine significant enrichment pathway. The phenylpropanoid biosynthesis, flavonoid biosynthesis, starch and sucrose metabolism, and plant hormone signal transduction were all significantly enriched in S3vs.S1 (**Fig. 3C**). Furthermore, in S4vs.S2, DEGs were significantly enriched to photosynthesis-antenna proteins, phenylpropanoid biosynthesis, ribosome, starch and sucrose metabolism, flavonoid biosynthesis (**Fig. 3D**). The phenylpropanoid biosynthesis (ko00940) pathway was enriched in five of the six comparison groups, with 58, 67, 61, 33, and 65 DEGs (**Fig. 3C, D, Supplementary Fig. S2**). Meanwhile, flavonoid biosynthesis, starch and sucrose metabolism, and plant hormone signal transduction also appeared in multiple groups such as S3vs.S1, S4vs.S1, S3vs.S2, and S4vs.S2.

### Global metabolomic profiles

Metabolic analysis showed that a total of 803 positive ion mode metabolites and 453 negative ion mode metabolites of hierarchical clustering analysis were identified in the 12 samples (**Fig. S12, S13**). PCA revealed a high degree of dispersion between each group (**Fig. 4A, B**), PC1 (37.13%) and PC2 (19.3%) separated POS and NEG metabolites, respectively. Most detected metabolites include lipids and lipid-like compounds (29.26% in POS, 35.32% in NEG) and phenylpropanoids and polyketides (17.85% in POS, 20.52% in NEG) (**Fig. 4C, D**). KEGG found 241 metabolites (**Supplementary Table S7**), mostly related to environmental, genetic, and metabolic processes. Hierarchical clustering analysis and PLS-DA model R2 and Q2 values for each comparison group were close to 1 (**Supplementary Fig. S6**), suggesting that the model is characterized by high stability and reliability. The heatmap illustrates twelve samples grouped into four primary clusters according to ion abundance and samples from each period were aggregated (**Fig. 4E**).

**Fig. 4.**
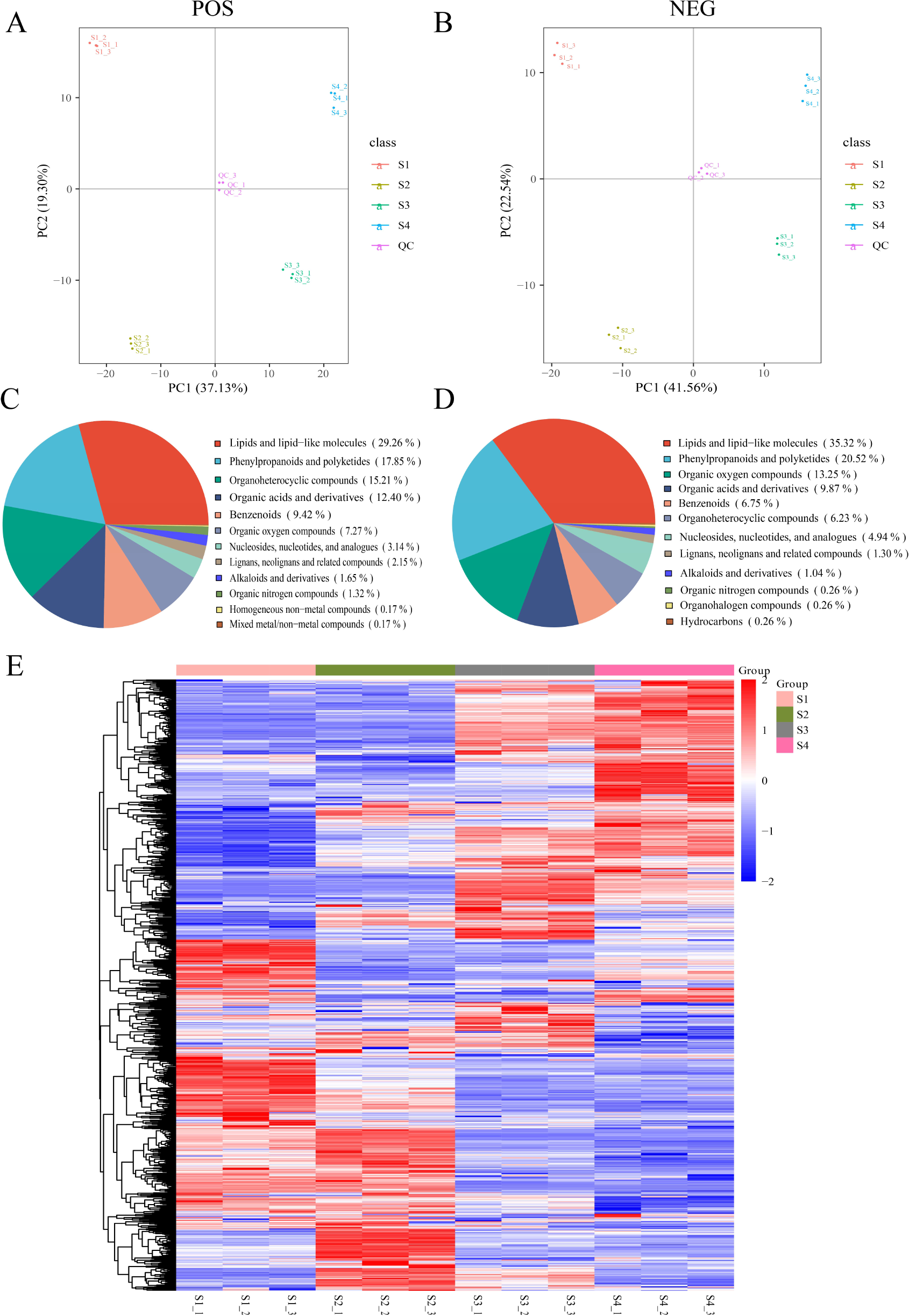
PCA of the metabolite profiles in POS (A) and NEG (B). Classification of metabolites in POS (C) and NEG (D). (E) Heatmap of total differential metabolite clustering in four different stages of bulbils. The expression levels are colored in red and blue for high and low expression.

### Identification of DEMs

Screening for differentially expressed metabolites was performed according to the criteria FC < 0.667 and p-value < 0.05 (**Supplementary Fig. S3, Table S6**). The comparison between S2vs.S1, S3vs.S1, and S4vs.S1 revealed the presence of 33 shared positive ionic DEMs (**Fig. 5A-D**). Prenol lipids (18.18%), carboxylic acids and derivatives (9.09%), coumarins and derivatives (6.06%), flavonoids, linear 1,3-diarylpropanoids, and purine nucleosides (6.06%) are among these metabolites. Fatty acyls (33.33%), undefined (16.67%), cinnamic acids and derivatives (5.56%), prenol lipids (5.56%), flavonoids (5.56%), and carboxylic acids and derivatives (5.56%) were among the 36 prevalent anionic DEMs. Five positive ion DEMs were found in bulbil developmentat stages (S3vs.S2, S4vs.S2, S4vs.S3): 2-[5-(ethylsulfonyl)-2-hydroxyanilino]carbonylbenzoic acid, cucurbitacin I, (5-L-Glutamyl)-L-amino acid, 1,7-bis(4-hydroxyphenyl)-5-methoxyheptan-3-one, and gibberellic acid (**Figure S8**). The top twenty up-regulated and down-regulated DEMs in S3vs.S1 and S4vs.S2 were both mainly lipids and lipid-like molecules and phenylpropanoids and polyketides (**Figure E, F**).

**Fig.5.**
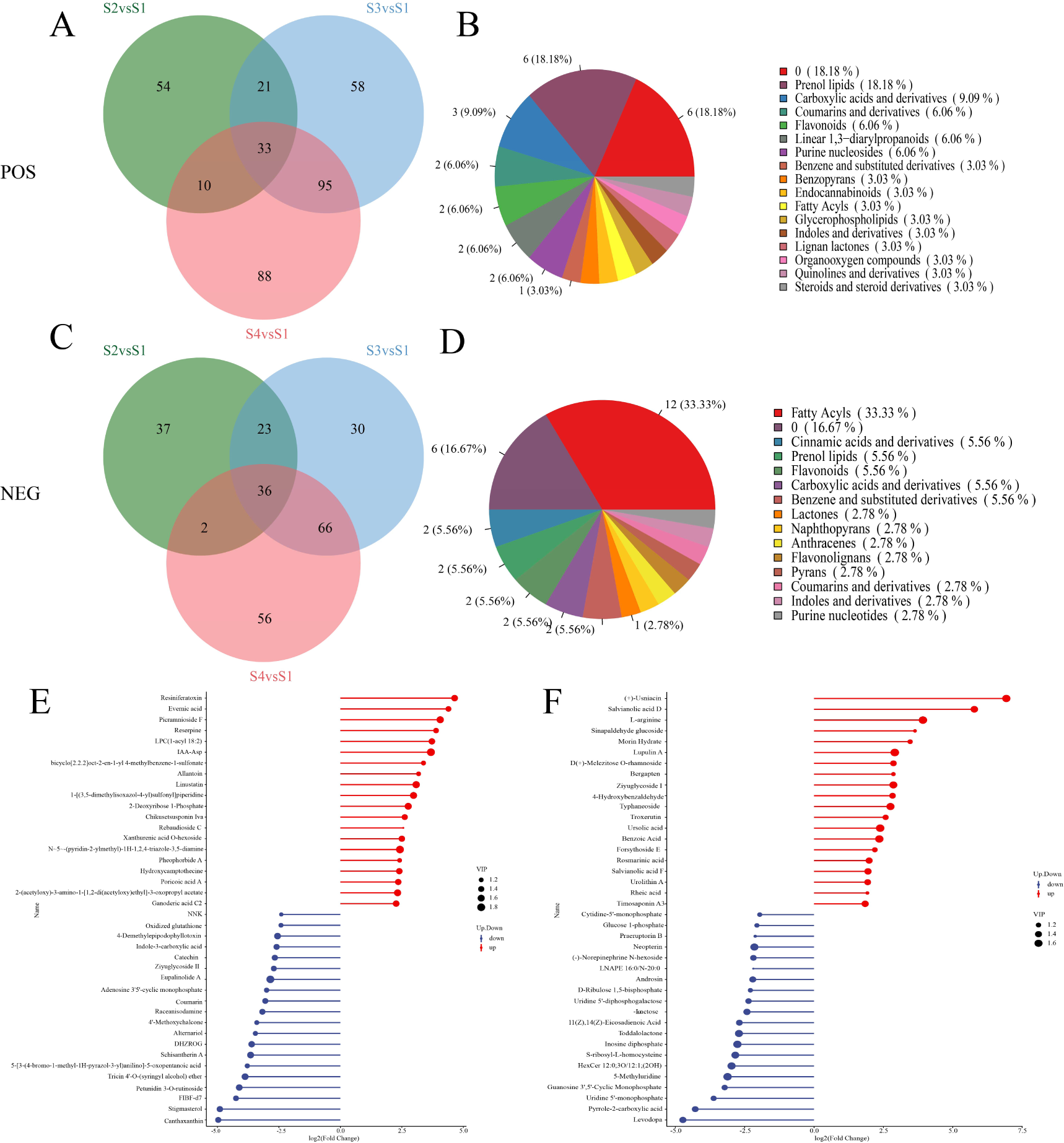
Venn diagram of DEMs shared among bulbil initiation in POS (A) and NEG (C). Classification of the shared DEMs in POS (B) and NEG (D). (E) The top 20 up and down-regulated DEMs in S3vs.S1. (F) The top 20 up and down-regulated DEMs in S4vs.S2.

### DEGs and DEMs involved in phytohormone signal transduction

The KEGG enrichment analysis of transcriptome revealed that 43 DEGs in total were assigned to plant hormone signal transduction (ko04075) pathways, including auxin (IAA), cytokinin (CK), ethylene, gibberellin (GA), abscisic acid (ABA), brassinosteroid (BR), jasmonic acid (JA), and salicylic acid (SA) signaling. The majority of these genes exhibited up-regulation in expression during S3 to S4, while showing down-regulation during S1 to S2 (**Fig. 6A**). Meanwhile, we detected ten metabolites related to plant hormone, which are mainly involved in the synthesis of auxin (IAA), gibberellin (GA), abscisic acid (ABA), jasmonic acid (JA) **(Fig. 6B**).

**Fig. 6.**
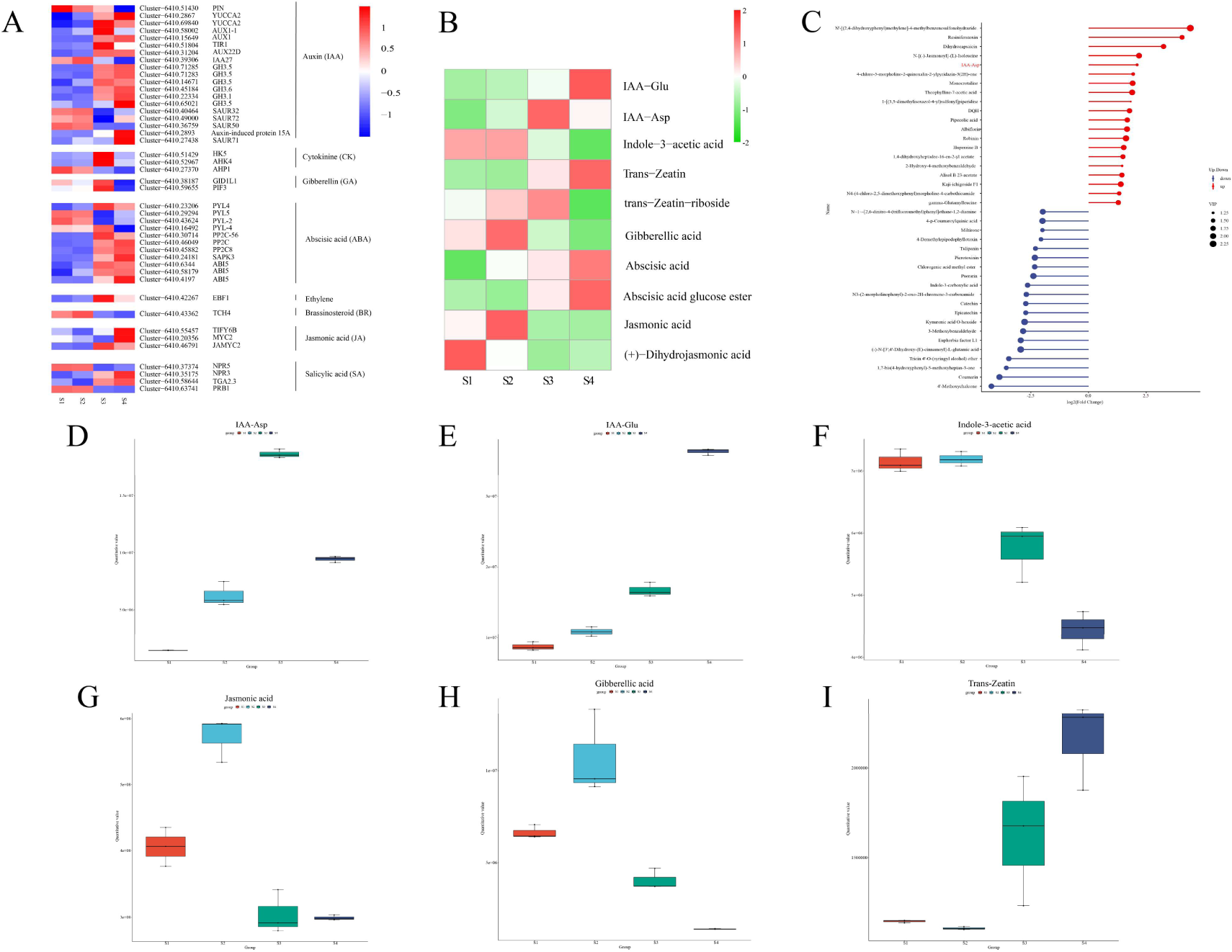
(A) Heatmap of genes involved in plant hormone signal transduction pathways. The expression levels are colored in red and blue for high and low expression. (B) Heatmap of metabolites involved in plant hormone signal transduction pathways. The expression levels are colored in red and green for high and low expression. (C) Stem plot of DEMs of S2vs.S1 in POS. (C) Box plot of IAA-Asp. (E) Box plot of IAA-Glu. (F) Box plot of Indole-3-acetic acid. (G) Box plot of jasmonic acid. (H) Box plot of gibberellic acid. (I) Box plot of Trans-Zeatin.

The expression profiles of nineteen differentially expressed genes in the IAA signal transduction pathway were analyzed (**Fig. 6A**). Many genes were up-regulated in S3 and S4 but down-regulated in S1 and S2. From S1 to S4, auxin influx carriers *PIN* were down-regulated. *YUCCA2* gene expression peaked in stages S4 and S3 of bulbil development. *AUX1-1* and *TIR1* had comparable expression patterns with peaks in stage S3. Five *GH3* genes (Cluster-6410.71285, Cluster-6410.71283, Cluster-6410.45184, Cluster-6410.22334, Cluster-6410.65021) and the auxin transporter protein 1 (AUX1) gene were up-regulated in S4 compared to S3. *AUX22D* gene was down-regulated in S1 and S2 and up-regulated in S3 and S4. *IAA27*, which encodes the auxin-responsive protein, was expressed from S1 to S2 but decreased from S3 to S4. The expression levels of three *SAUR* genes (*SAUR32*, *SAUR72*, *SAUR50*) were likewise higher in S1 and S2 than S3 and S4. IAA-Asp from organic acids and derivatives was an up-regulated DEM in stage S3 (**Fig. 6C, D**), which is correlated with the expression pattern of the *TIR1* gene. However, IAA-Glu showed an up-regulated trend in S4 (**Fig. 6E**), and Indole-3-acetic acid was highly expressed in S1 and S2 where bulbils were in the initiation stages (**Fig. 6B, F**).

In the CK signal transduction pathway, the *AHK* gene was highly expressed in S3 and the expression level of the *AHP1* gene was the highest in S1. *PIF3* gene and gibberellic acid in the gibberellin signal transduction pathway exhibited the highest expression in S2 (**Fig. 6A, H**). Trans-Zeatin and Trans-Zeatin-riboside were detected in metabolomic analysis, both showing high expression in S3 (**Fig. 6B, I**). The ABA signaling system had four *PYL* genes, three *PP2C* genes, one *SAPK* gene, three *ABI5* genes, abscisic acid, and its glucose ester. The expression of abscisic acid and abscisic acid glucose ester were similar in S4 and S3 (**Fig. 6B**), which were highly expressed in S4 when bulbils were mature. In contrast, the *XTH* gene encoding the BR signal transduction pathway showed high expression in S1 and S2. In the JA signal transduction pathway, the *TIFY6B* gene and *MYC2* gene peaked during S4, and the *JAMYC2* gene showed the highest expression in S3, while jasmonic acid (**Fig. 6G**) and (+)-Dihydrojasmonic acid were highly expressed in S2 and S1. Four genes were identified in the SA signal transduction pathway, with the *NPR5* gene and *PRB1* gene showing high expression in S1 and S2, the *NPR3* gene, and the *TGA2.3* gene displaying up-regulated expression in S4 vs S3.

### DEGs and DEMs Involved in starch and sucrose metabolism

KEGG analysis showed that 18 DEGs encoding enzymes related to starch and sucrose metabolism were identified (**Fig. 7**). 53, 64, 55, 64, and 20 DEGs between S3vs.S1, S4vs.S1, S3vs.S2, S4vs.S2 and S4vs.S3, respectively, were enriched. The *INV* gene is a crucial enzyme involved in the hydrolysis of sucrose. Notably, three genes encoding *INV* consistently demonstrated high expression levels from S1 to S3. The expression levels of starch synthase genes, including the *SUS2* gene and *SSS* gene, were significantly elevated in stages S1 and S2. Glucose 1-phosphate was highly expressed in S1 and S2 compared to S3 and S4 (**Fig. 7B**) and this is probably correlated to the expression of *INV* gene and *SUS2* gene. *ENPP1_3* gene was up-regulated in stage S3, while *AGPS* gene showed increased expression in stages S3 and S4. *SBE* gene expression increased gradually throughout the stages, peaking in stage S4. *AMY* gene and *malQ* gene showed increasing pattern in S4vs.S3, which may be related to starch breakdown as bulbils grow and mature. We found that S3 had the highest expression of *TPS* gene, which regulates axillary bud development through sugar signaling by generating trehalose-6-phosphate. The *GLU* gene was also expressed at higher levels in S1 and S2, increasing cellodextrin and cellobiose synthesis. In contrast, the *BGLU* gene was up-regulated in S3 and S4, increasing D-Glucose synthesis.

**Fig. 7.**
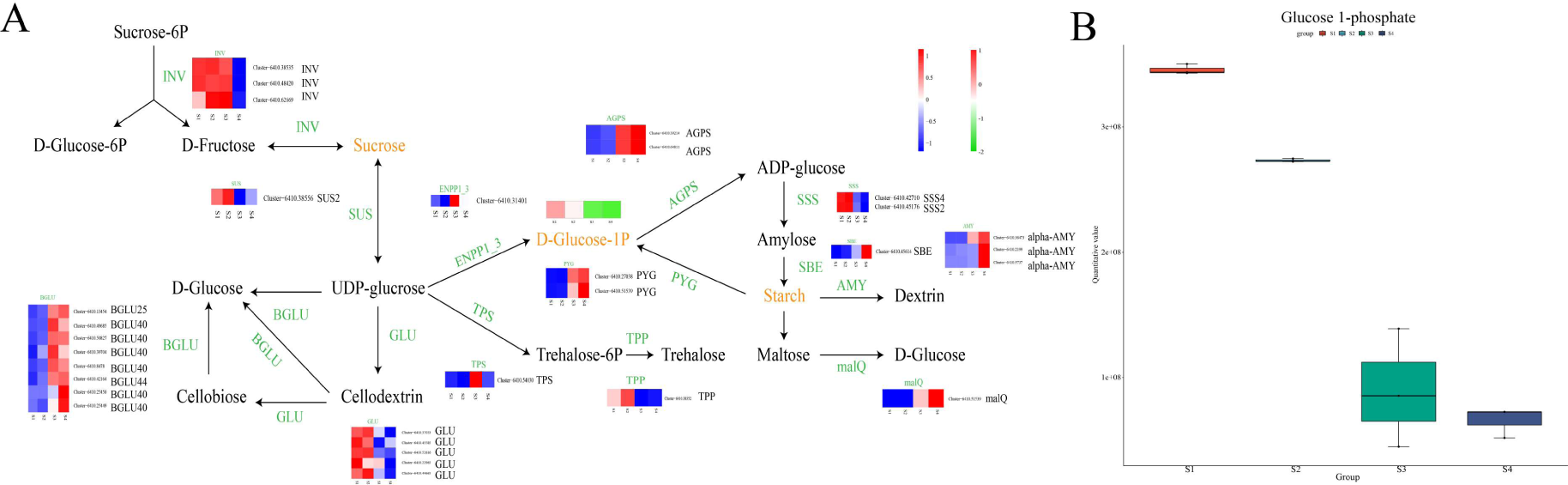
(A) Heatmap of genes and metabolites involved in starch and sucrose metabolism pathway. The expression levels are colored in red and blue, red and green, for high and low expression. (B) Box plot of Glucose 1-phosphate.

### DEGs and DEMs Involved in phenylpropanoid biosynthesis and flavonoid biosynthesis

Phenylpropanoid biosynthesis is an upstream pathway for flavonoid biosynthesis. We identified 49 DEGs and 9 DEMs in phenylpropanoid biosynthesis and flavonoid biosynthesis (**Fig. 8A, B**). Three *PAL* genes (Cluster-6410.27707, Cluster-6410.40307, Cluster-6410.24298) and *CYP73A* gene were highly expressed in S1, which is relative to the high expression of p-coumaric acid and coumarin in S1 (**Fig. 8C, D**). During the four stages, the *bgIB* gene increased. In S2, three *COMT* and four *4CL* genes were up-regulated. Nine *HCT* genes are highly expressed in S4. S1 and S2 had higher *CYP98*, *CCoAOMT*, and *CCR* gene expression than S3 and S4. Two *DFR* genes were strongly expressed in S3 and S4, but the other two were in S1 and S2. The expression of three *CHS* genes rose from S1 to S4 and peaked in S4. On the contrary, *CYP75A* gene was down-regulated during the four stages. Butein, naringenin chalcone, and myricetin were down-regulated from S1 to S4 and showed the highest expression in S1 (**Fig. 8B**). However, scopoletin showed an upward expression and peaked in S4 (**Fig. 8H**). In addition, periodical showed the highest expression in S3, and (+)-catechin showed higher expression in S1 and S4 (**Fig. 8G, E**).

**Fig. 8.**
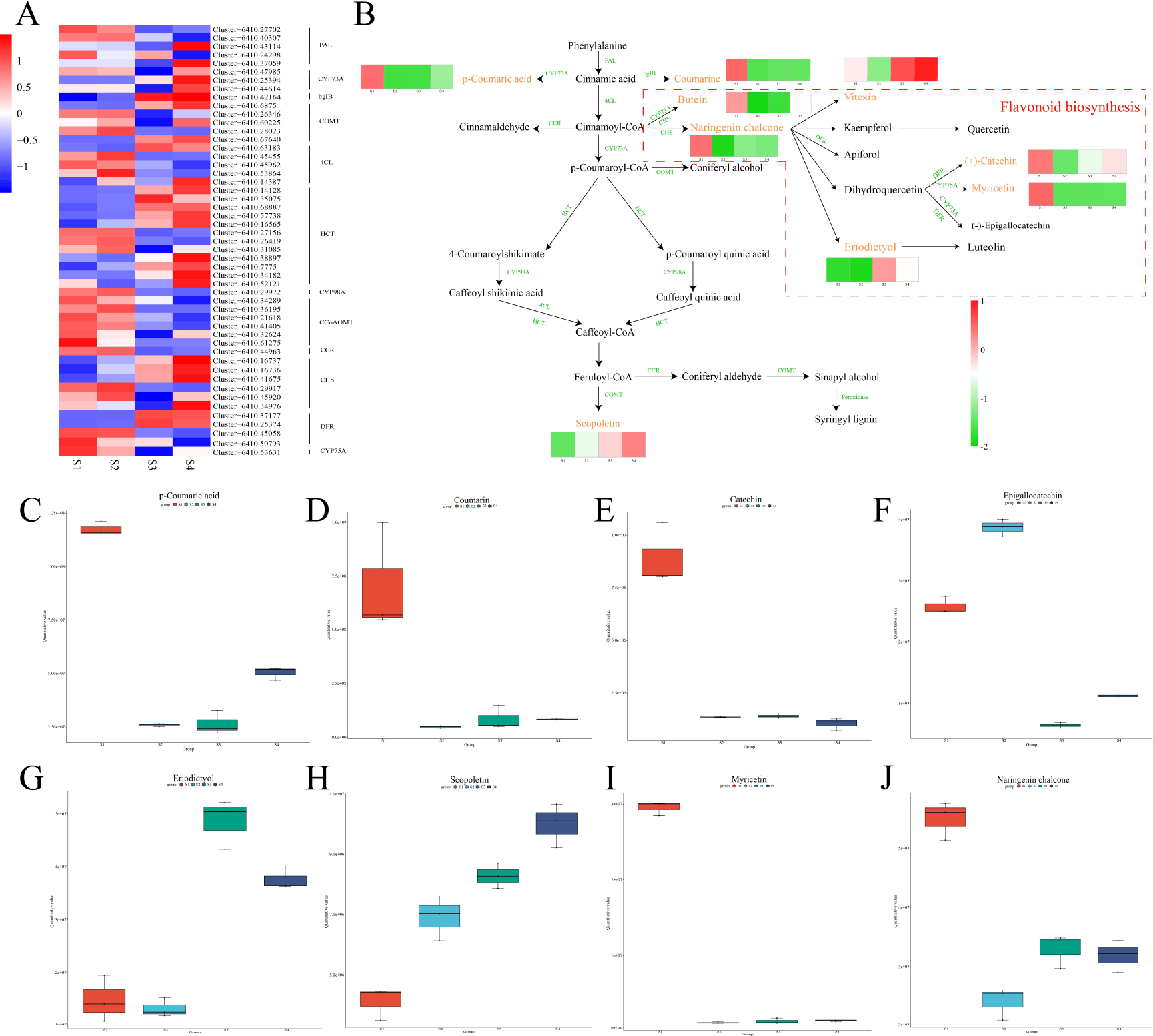
(A, B) Heatmap of genes and metabolites involved in phenylpropanoid biosynthesis and flavonoid biosynthesis pathways. The expression levels are colored in red and blue, red and green for high and low expression. (C) Box plot of p-coumaric acid. (D) Box plot of coumarin. (E) Box plot of catechin. (F) Box plot of epigallocatechin. (G) Box plot of eriodicyol. (H) Box plot of scopoletin. (I) Box plot of myricetin. (I) Box plot of naringenin chalcone.

### Correlation analysis between the transcriptome and metabolome in key KEGG pathways

Six sets of samples were compared, with S2vs.S1 having the fewest differential expreesed genes and metabolites while S4vs.S1 having the most. Samples with similar phases had fewer differences than those with farther apart stages (**Supplementary Fig. 16**). Phenylpropanoid biosynthesis, flavonoid biosynthesis, ABC transporters, galactose metabolism, pentose phosphate pathway, and plant hormone signal transduction are co-enriched in the transcriptome and metabolome during bulbil initiation (S3vs.S1, S4vs.S1). Flavonoid biosynthesis, plant hormone signal transduction, and ABC transporters pathways involved in gene and metabolite expression increased throughout bulbil growth and development (S3vs.S2, S4vs.S2, S4vsS3) (**Supplementary Table S9)**. Genes and metabolites of starch and sucrose metabolism pathway in S4vs.S2 and amino sugar and nucleotide sugar metabolism pathway in S3vs.S2 were significantly expressed compared to other comparison groups (**Figure 9A-D**). Differential metabolites and genes were mapped to iPath, lipid metabolism, amino acid metabolism, energy metabolism, and biosynthesis of other secondary metabolites pathways for bulbil initiation stages. Bulbil growth and development have been associated with more pathways, such as carbohydrate metabolism, xenobiotics biodegradation and metabolism, nucleotide metabolism, metabolism of cofactors and vitamins, and metabolism of other amino acids (**Supplementary Fig. S10**).

**Fig. 9.**
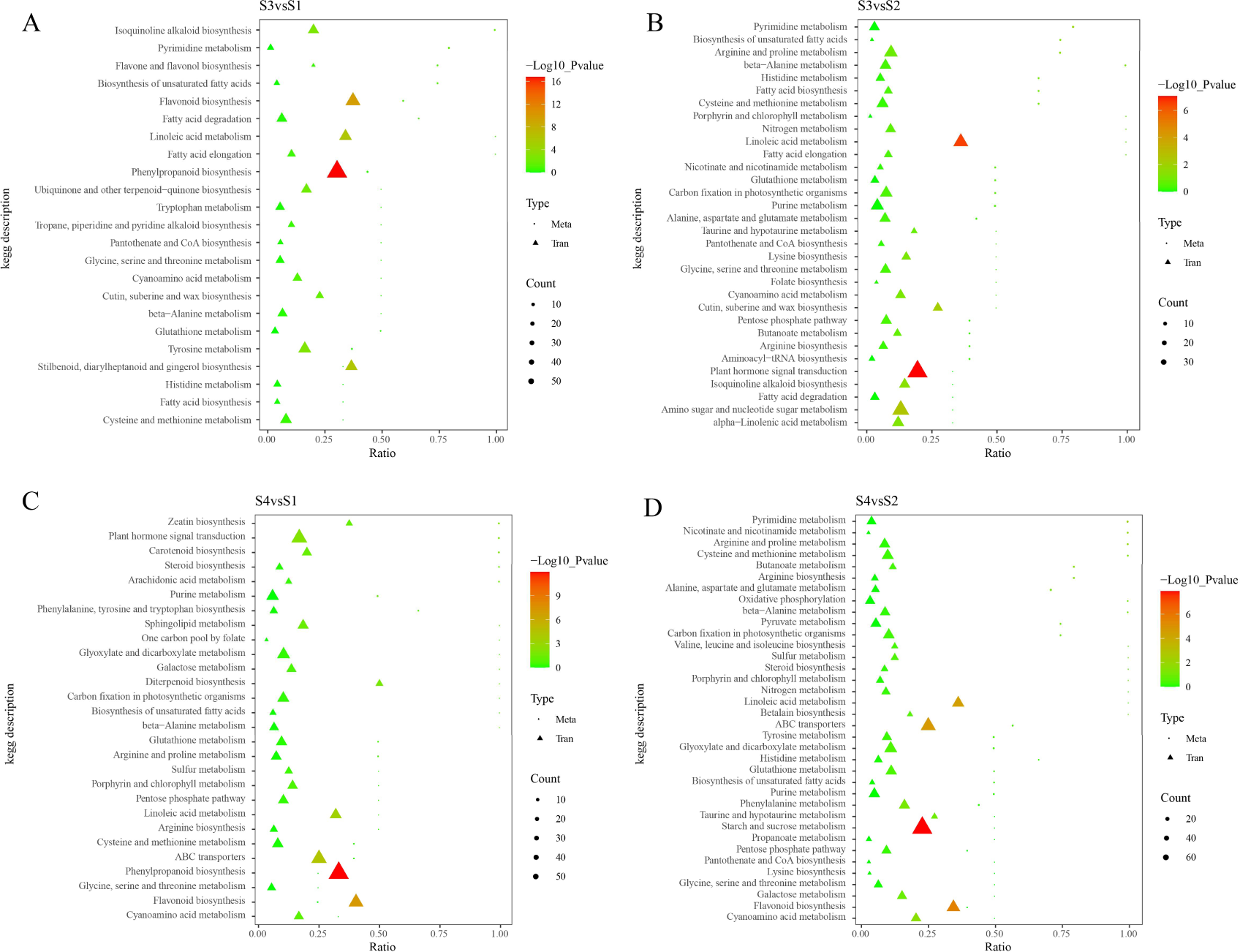
The shared KEGG pathway scatterplot of DEGs and DEMs in S3vs.S1 (A), S4vs.S1 (B), S3vs.S2 (C), S4vs.S2 (D).

## Discussion

### The formation of bulbil in the endangered fern *Cystopteris chinensis* is a unique phenomenon

Known fern species that can produce bulbil are currently found on *Woodwardia orientalis* (Baskaran, 2016), *Woodwardia unigemmata*, *Athyrium Clarke*, *Monachosorum henryi* (Luo *et al*., 2014), and *Cystopteris bulbifera* (Moran, 2004) and so on. The bulbil of *C. chinensis* shows an annual type, forming from May to July each year. We found that the bulbils are inserted on the lateral veins at the tip of the pinnae and the color of *C. chinensis* bulbil gradually changes from a near-transparent light green to a darker green during development, with a brownish change and adventitious-root-like formation at the apical and lower ends of the bulbil, which may be related to the accumulation of pigments, amyloplasts, and flavonoids. Under transmission electron microscopy, it was observed that during the formation, differentiation, and maturation of *C. chinensis* bulbil, there was a continuous increase in cell volume. Both starch granules and pigment granules showed a consistent increase. The development of vascular bundles facilitated nutrients such as sucrose to transport from the fronds to bulbil, contributing to the overall expansion of bulbil. These results suggest the origin of bulbil in *C. chinensis* may have been attributed to the organogenesis of meristematic cells at the tip of the pinnule where bulbils attach (Garcês *et al*., 2007).

### Phytohormones control the initiation and development of bulbil in *C. chinensis*

Plant hormone is important to the growth and development of plants (Santner *et al*., 2009). We found genes and metabolites related to phytohormones that influence bulbil initiation and development. Auxin is involved in bulbils initiation and plays a role in bulbils development according to our results. Indole-3-acetic acid (IAA) was highly expressed in S1 and S2. In S3 and S4, IAA-Asp and IAA-Glu, the storage forms of IAA, were upregulated. We found the auxin transport gene *PIN* was significantly expressed in S1 and gradually down-regulated to S4, suggesting auxin efflux decreased (Křeček *et al*., 2009). The Arabidopsis mutant *yuc1d* exhibits an auxin overproduction phenotype, indicating YUC family is related to the synthesis of IAA (Zhao, 2010). The high expression of Two *YUCCA2* genes from S3 to S4 can be associated with the accumulation of IAA-Glu and IAA-Asp during bulbil developmental stages. Interestingly, the *IAA27* gene in the auxin signaling showed the highest expression in S2 where bulbils appeared at initiation stages, indicating that auxin is involved in the initiation of bulbils. The small auxin-up RNA (*SAUR*) gene can control auxin synthesis and signaling, we found *SAUR32*, *SAUR72,* and *SAUR50* were highly expressed in S2 but down-regulated in the bulbil developmental stages except *SAUR71* peaked at S4, suggesting *SAUR* genes in *C. chinensis* attribute to bulbils initiation. It is also found in *L. radiata* that *SAUR50/61* genes are up-regulated when bulbil initiation occurs (Xu *et al*., 2020). *GH3* genes are involved in catalyzing IAA conjugation to amino acids in arabidopsis and can reduce IAA concentration (Luo *et al*., 2023). We found that most *GH3* genes were up-regulated in S3 and S4 and peaked at S4, which indicates that IAA concentration is high in bulbil initiation stages but degrades at the same time as bulbil enters mature stage S4. In conclusion, auxin plays a crucial role in triggering bulbil initiation during the S2 stage. However, auxin transport becomes restricted at a later stage as bulbils approach maturity in S4.

Cytokine can stimulate bud outgrowth and is influenced by IAA (Ferguson and Beveridge, 2009). We found CK receptors *HK5* gene and *AHK4* gene were significantly up-regulated in S3, while in *Asiatic hybrid* lily, *AHK2* is significantly up-regulated during bulbil initiation (Liang *et al*., 2023). Most of the *AHPs* are positive regulators of cytokinin signaling and affect multiple aspects of plant development (Hutchison *et al*., 2006), our results showed *AHP1* gene was highly expressed in S1 and S2. However, trans-zeatin-riboside and trans-zeatin as precursor substances of CK synthesis, showed high expression in S2, S3, and S4, respectively. These results indicate that cytokinine is responsible for bulbils initiation and differentiation. Other genes and metabolites related to phytohormones including BR, GA, SA, ABA, ETH, and JA showed different expressions. *NPR5* gene, and *PRB1* gene in salicylic acid synthesis were up-regulated in S1 and S2, which is similar to changes in *L. radiata* (Xu *et al*., 2020) indicating salicylic acid promotes bulbils initiation. *GID1L*, *PIF3* gene in gibberellin synthesis and *EBF1* in ethylene synthesis both reached peak expression in S3, meanwhile gibberellic acid showed high expression in S3 and S4, suggesting gibberellin and ethylene together affect bulbils differentiation. *PYL* gene was reported to be down-regulated in bulbil initiation and can be significantly up-regulated after treatment with PCX (Guo *et al*., 2022).

### Starch and sucrose biosynthesis plays a crucial role in the initiation and development of bulbil *in C. chinensis*

The secret behind bulbil production in plants is mainly ascribed to the starch and sucrose changes during bulbil formation, many studies have proved that soluble sugar promotes the initiation of bulbil and starch promotes the development of bulbil (Yang *et al*., 2017; Wu *et al*., 2020; Xu *et al*., 2020; Cao *et al*., 2023; Shu *et al*., 2024). Bulbil, as a kind of sink in plants, requires sucrose to be degraded into hexoses or their derivates for biosynthetic processes (Ruan *et al*., 2010). *INV* gene hydrolyses sucrose into glucose and fructose and functions in primary carbon metabolism (Barratt *et al*., 2009). In our study, three *INV* genes were highly expressed during the initiation and differentiation stages of bulbil, indicating glucose and fructose regulated the initiation and early development of bulbil. We also detected the *SUS2* gene which degrades sucrose into UDP-glucose showing the highest expression in S2. In addition, glucose 1-phosphate, which is a key intermediate in several major carbon fluxes, such as starch, sucrose, and cellulose biosynthesis (Fettke *et al*., 2011), showed high expression in S1 and S2 consistent with the expression of *SSS* genes. These results suggest sucrose synthesis plays a key role in the initiation of bulbil. T6P is responsible for plant growth and development leading to branching phenotypes, it was found that embryo-lethal *A. thaliana* tps1 mutants are rescued by the expression of *E. coli TPS* (Schluepmann *et al*., 2003). *TPS* gene and *TPP* gene in *C. chinensis* were up-regulated in S3 and S2 respectively, which suggests sucrose signaling controls the early development of bulbils. We observed genes related to starch synthesis including *AGPs*, *SBE*, *PYG,* and *AMY* were highly expressed in S3 and S4 and all these genes peaked at S4, indicating that starch plays a major role in the development of bulbil, especially during maturation.

### Phenylpropanoid biosynthesis and flavonoid biosynthesis regulate the pigment and vascular bundle formation of bulbils in *C. chinensis*

Phenylpropanoid metabolism is highly adaptable to different stages of plant development and changing environmental conditions. This pathway produces a variety of products that play important roles in plant growth and interaction with the environment. Flavonoids constitute another important group of phenolic compounds synthesized via the phenylpropanoid pathway (Dong and Lin, 2021). Phenylpropanoid biosynthesis is particularly significant in the formation of lignin, a complex polymer that reinforces cell walls in vascular tissues (Zhao, 2016). In our study, two *PAL* genes showed peak expression in S4, which is consistent with the vascular bundle formation in bulbil maturation stages. P-coumaric acid and coumarins generated from cinnamic acid were up-regulated in S1 when bulbils had not occurred. Their expressions are negatively correlated with *CYP73A* and *bgIB*, which are the genes controlling the synthesis. 4-Coumarate-CoA ligase catalyzes the third step of the general phenylpropanoid pathway, the formation of p-coumaroyl-CoA in an adenosine triphosphate-dependent manner (Gui *et al*., 2011). We observed that 4 of 5 *4CL* genes were highly expressed in S2, suggesting those genes play a crucial role during bulbil initiation. Silencing *HCT* downregulated lignin biosynthesis, directing metabolic pathways towards flavonoid production, highlighting HCT’s crucial role in lignin biosynthesis (Hoffmann *et al*., 2004). 10 *HCT* genes and 4 *COMT* genes showed an up expression during bulbil development, which is consistent with the expression of scopoletin from S3 to S4, indicating lignin in bulbil was increased possibly to form a vascular bundle that transports nutrients. CHS plays a critical role as a key enzyme in the biosynthetic pathway responsible for secondary flavonoid metabolites, specifically contributing to the accumulation of flavonols and overexpression of flavonoid structural genes such as *CHS* and *DFR* to increase the levels of flavonol glycosides and anthocyanins (Cui *et al*., 2014). As bulbil developed, we observed pigment formation under TEM. This is correlated to the high expression of *CHS* genes in S2 and S4. However, butein, naringenin chalcone, (+)-catechin, and myricetin all showed specifically high expression in S1 instead of eriodictyol peaking in S3, suggesting the accumulation of flavonols through the initiation and development of bulbil. In addition, the *CYP75A* gene was gradually down-regulated from S1 to S4, which is positively consistent with the expression of myricetin.

## Conclusion

*Cystopteris chinensis* (Ching) X.C. Zhang & R. Wei is nearly extinct due to poor spore reproduction. It uses bulbil propagation to make new plants to maintain its population. We used morphological observations, transmission electron microscopy to study ultrastructures, and transcriptome and metabolomic investigations of bulbil at different developmental stages to investigate the molecular processes of this unusual propagation. **Figure 10** shows a molecular and genetic regulatory network model for C. chinensis bulbil initiation and development. We discovered that plant hormones, starch, sucrose, phenylpropanoids, and flavonoid production are key to bulbil formation and growth. Many plant hormones are involved especially auxin and cytokinin. Auxin was high during bulbil production but decreased in the four stages and mostly existed as IAA-Glu and IAA-Asp. *GH3* genes are significantly expressed during bulbil maturation, causing auxin degradation late in development. Cytokinin also helps bulbil growth begin. In bulbil ultrastructural cells, starch granules increased during development, while transcriptome data showed that genes that regulate glucose and fructose, such as *INV* and *SUS*, and metabolites like glucose 1-phosphate were highly expressed in bulbil initiation stages. The *TPS* gene converted *T6P* into starch during bulbil differentiation, while genes like *AGPs*, *SBE*, *PYG*, and *AMY* were highly expressed during bulbil maturation, adding to starch buildup. We found that phenylpropanoid and flavonoid production controlled bulbil pigment and vascular bundle development in *C. chinensis*. These results describe a basic molecular mechanism and metabolite accumulation of bulbil initiation and growth. Understanding how these candidate genes govern bulbils will help *C. chinensis* reproduce and conserve.

**Fig. 10.**
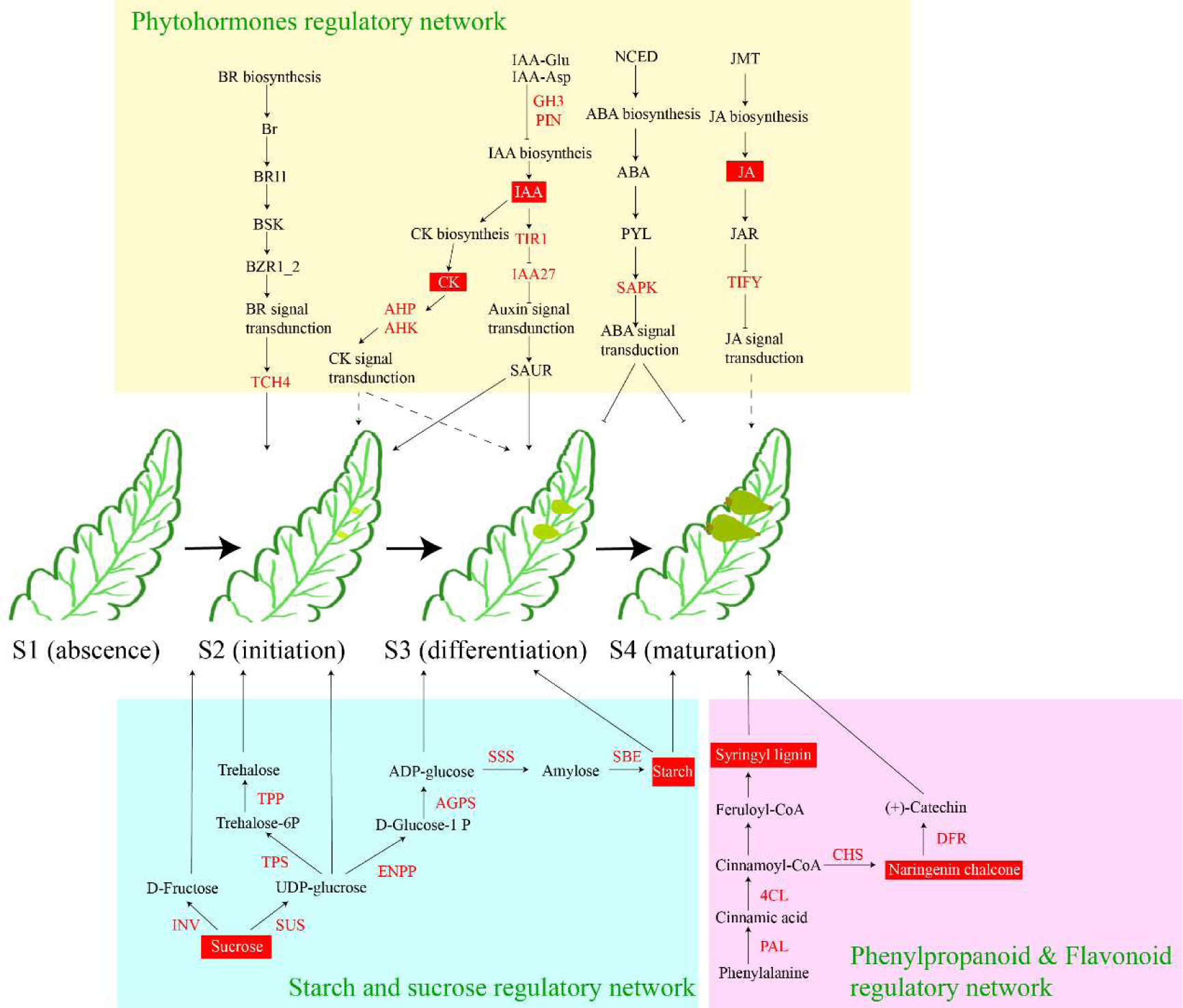
A proposed model to explain the molecular mechanism of bulbils initiation and development in *C. chinensis.* The red words refer to key genes.

## Acknowledgments

This study was supported by Natural Science Foundation of Sichuan Province: Study on the Reproductive Characteristics and Formation Mechanism of *Cystoperis chinensis,* a National-level Protected Plant (no.22NSFSC0230) in China).

## Author contributions

Xiaohong Chen conceived the program, An Yu and Wenkai Xi designed the experiment, and An Yu performed most of the experiments, with assistance from Xia Zhao, Yazhu Wang, Zhihong Gong and Xiaofeng Zhou. An Yu analyzed the transcriptome data and dealt with the figures. The manuscript was written by An Yu. Xiaohong Chen and Wenkai Xi modified the manuscript. All authors read and approved the final manuscript.

## Notes

### Competing Interest Statement

The authors have declared no competing interest.

## References

Abraham-Juárez MJ, Martínez-Hernández A, Leyva-González MA, Herrera-Estrella L, Simpson J. 2010. Class I KNOX genes are associated with organogenesis during bulbil formation in Agave tequilana. Journal of Experimental Botany 61, 4055–4067.

Arizaga S, Ezcurra E. 2002. Propagation mechanisms in Agave macroacantha (Agavaceae), a tropical arid-land succulent rosette. American Journal of Botany 89, 632–641.

Barratt DHP, Derbyshire P, Findlay K, Pike M, Wellner N, Lunn J, Feil R, Simpson C, Maule AJ, Smith AM. 2009. Normal growth of Arabidopsis requires cytosolic invertase but not sucrose synthase. Proceedings of the National Academy of Sciences of the United States of America 106, 13124–13129.

BASKARAN XR. 2016. First insights on next-generation sequencing and biochemical analysis of an apomictic fern for vivipary character. postdoc, Guangzhou: Sun Yat-Sen University.

Cai Y, Yu L, Wen Z, Gong W, Yang H, Zhao C. 2016. A Study of the Method of Isolated Culture of Cystoathyrium chinensis Spores. Journal of Sichuan Forestry Science and Technology 37, 76–79.

Cao T, Wang S, Ali A, et al. 2023. Transcriptome and metabolome analysis reveals the potential mechanism of tuber dynamic development in yam (Dioscorea polystachya Turcz.). LWT 181, 114764.

Chen H, Yu L, Wen Z, Wang D, Wu S. 2016. Propagation Technique of Cystoathyrium chinese Spores. Journal of Sichuan Forestry Science and Technology 37, 97–99.

Ching RC. 1966. Three new fern genera. Acta Phytotaxonomica Sinica 11, 17–30.

Cui L-G, Shan J-X, Shi M, Gao J-P, Lin H-X. 2014. The miR156-SPL9-DFR pathway coordinates the relationship between development and abiotic stress tolerance in plants. The Plant Journal: For Cell and Molecular Biology 80, 1108–1117.

D. Bell A, Bryan A. 2008. Plant Form: An Illustrated Guide to Flowering Plant Morphology. Portland, OR: Timber Press.

Dong N-Q, Lin H-X. 2021. Contribution of phenylpropanoid metabolism to plant development and plant-environment interactions. Journal of Integrative Plant Biology 63, 180–209.

Engel A, Colliex C. 1993. Application of scanning transmission electron microscopy to the study of biological structure. Current Opinion in Biotechnology 4, 403–411.

Ferguson BJ, Beveridge CA. 2009. Roles for Auxin, Cytokinin, and Strigolactone in Regulating Shoot Branching. Plant Physiology 149, 1929–1944.

Fettke J, Malinova I, Albrecht T, Hejazi M, Steup M. 2011. Glucose-1-Phosphate Transport into Protoplasts and Chloroplasts from Leaves of Arabidopsis. Plant Physiology 155, 1723– 1734.

Garcês HMP, Champagne CEM, Townsley BT, Park S, Malhó R, Pedroso MC, Harada JJ, Sinha NR. 2007. Evolution of asexual reproduction in leaves of the genus *Kalanchoë*. Proceedings of the National Academy of Sciences 104, 15578–15583.

Grabherr MG, Haas BJ, Yassour M, et al. 2011. Full-length transcriptome assembly from RNA-Seq data without a reference genome. Nature Biotechnology 29, 644–652.

Gui J, Shen J, Li L. 2011. Functional characterization of evolutionarily divergent 4-coumarate:coenzyme a ligases in rice. Plant Physiology 157, 574–586.

Guo C, Li J, Li M, Xu X, Chen Y, Chu J, Yao X. 2022. Regulation Mechanism of Exogenous Brassinolide on Bulbil Formation and Development in Pinellia ternata. Frontiers in Plant Science 12.

Haufler CH, Pryer KM, Schuettpelz E, Sessa EB, Farrar DR, Moran R, Schneller JJ, Watkins JE Jr, Windham MD. 2016. Sex and the Single Gametophyte: Revising the Homosporous Vascular Plant Life Cycle in Light of Contemporary Research. BioScience 66, 928–937.

Higuchi H, Amaki W. 1989. Effects of 6-benzylaminopurine on the organogenesis of Asplenium nidus L. through in vitro propagation. Scientia Horticulturae 37, 351–359.

Higuchi H, Amaki W, Suzuki S. 1987. In vitro propagation of *Nephrolepis cordifolia* Prsel. Scientia Horticulturae 32, 105–113.

Hoffmann L, Besseau S, Geoffroy P, Ritzenthaler C, Meyer D, Lapierre C, Pollet B, Legrand M. 2004. Silencing of Hydroxycinnamoyl-Coenzyme A Shikimate/Quinate Hydroxycinnamoyltransferase Affects Phenylpropanoid Biosynthesis[W]. The Plant Cell 16, 1446–1465.

Hutchison CE, Li J, Argueso C, et al. 2006. The Arabidopsis Histidine Phosphotransfer Proteins Are Redundant Positive Regulators of Cytokinin Signaling. The Plant Cell 18, 3073– 3087.

Kenrick P, Crane P. 1997. The origin and early diversification of land plants: A cladistic study. Washington: Smithsonian.

Křeček P, Skůpa P, Libus J, Naramoto S, Tejos R, Friml J, Zažímalová E. 2009. The PIN-FORMED (PIN) protein family of auxin transporters. Genome Biology 10, 249.

León P, Sheen J. 2003. Sugar and hormone connections. Trends in Plant Science 8, 110–116.

Letunic I, Yamada T, Kanehisa M, Bork P. 2008. iPath: interactive exploration of biochemical pathways and networks. Trends in Biochemical Sciences 33, 101–103.

Li F-W, Brouwer P, Carretero-Paulet L, et al. 2018. Fern genomes elucidate land plant evolution and cyanobacterial symbioses. Nature Plants 4, 460–472.

Li T, Li SQ, Luo R. 2012. Morphological observation and anatomical study on bulbil development of Lilium sulphureum. Acta Botanica Boreali-Occidentalia Sinica 32, 85–89.

Li X, Wang C, Cheng J, Zhang J, da Silva JAT, Liu X, Duan X, Li T, Sun H. 2014. Transcriptome analysis of carbohydrate metabolism during bulblet formation and development in Lilium davidii var. unicolor. BMC Plant Biology 14, 358.

Liang J, Chen Y, Hou J, et al. 2023. Cytokinins influence bulblet formation by modulating sugar metabolism and endogenous hormones in Asiatic hybrid lily. Ornamental Plant Research 3.

Luo R, Du YS, Sun YY, Cao ZJ. 2014. Morphological Observation and Anatomical Study on Bulbil Development of Pinellia ternata. Acta Botanica Boreali-Occidentalia Sinica 34, 1776– 1781.

Luo P, Li T-T, Shi W-M, Ma Q, Di D-W. 2023. The Roles of GRETCHEN HAGEN3 (GH3)-Dependent Auxin Conjugation in the Regulation of Plant Development and Stress Adaptation. Plants 12, 4111.

Moran RC. 2004. A Natural History of Ferns.

Pryer KM, Schneider H, Smith AR, Cranfill R, Wolf PG, Hunt JS, Sipes SD. 2001. Horsetails and ferns are a monophyletic group and the closest living relatives to seed plants. Nature 409, 618–622.

Ranker TA, Haufler CH. 2008. Biology and Evolution of Ferns and Lycophytes. Cambridge, UK; New York: Cambridge University Press.

Ruan Y-L, Jin Y, Yang Y-J, Li G-J, Boyer JS. 2010. Sugar input, metabolism, and signaling mediated by invertase: roles in development, yield potential, and response to drought and heat. Molecular Plant 3, 942–955.

Santner A, Calderon-Villalobos LIA, Estelle M. 2009. Plant hormones are versatile chemical regulators of plant growth. Nature Chemical Biology 5, 301–307.

Schluepmann H, Pellny T, van Dijken A, Smeekens S, Paul M. 2003. Trehalose 6-phosphate is indispensable for carbohydrate utilization and growth in Arabidopsis thaliana. Proceedings of the National Academy of Sciences of the United States of America 100, 6849–6854.

Schneider H, Schuettpelz E, Pryer KM, Cranfill R, Magallón S, Lupia R. 2004. Ferns diversified in the shadow of angiosperms. Nature 428, 553–557.

Schrimpe-Rutledge AC, Codreanu SG, Sherrod SD, McLean JA. 2016. Untargeted Metabolomics Strategies—Challenges and Emerging Directions. Journal of the American Society for Mass Spectrometry 27, 1897–1905.

Shu F, Wang D, Sarsaiya S, Jin L, Liu K, Zhao M, Wang X, Yao Z, Chen G, Chen J. 2024. Bulbil initiation: a comprehensive review on resources, development, and utilisation, with emphasis on molecular mechanisms, advanced technologies, and future prospects. Frontiers in Plant Science 15.

Thomas BA, Cleal CJ. 2022. Pteridophytes as primary colonisers after catastrophic events through geological time and in recent history. Palaeobiodiversity and Palaeoenvironments 102, 59–71.

Tizro P, Choi C, Khanlou N. 2019. Sample Preparation for Transmission Electron Microscopy. In: Yong WH, ed. Biobanking: Methods and Protocols. New York, NY: Springer New York, 417– 424.

Walck JL, Cofer MS, Hidayati SN. 2010. Understanding the germination of bulbils from an ecological perspective: a case study on Chinese yam (Dioscorea polystachya). Annals of Botany 106, 945–955.

Wang C-N, Cronk QCB. 2003. Meristem fate and bulbil formation in Titanotrichum (Gesneriaceae). American Journal of Botany 90, 1696–1707.

Wang C-N, Möller M, Cronk QCB. 2004. Altered expression of GFLO, the Gesneriaceae homologue of FLORICAULA/LEAFY, is associated with the transition to bulbil formation in Titanotrichum oldhamii. Development Genes and Evolution 214, 122–127.

Want EJ, Masson P, Michopoulos F, Wilson ID, Theodoridis G, Plumb RS, Shockcor J, Loftus N, Holmes E, Nicholson JK. 2013. Global metabolic profiling of animal and human tissues via UPLC-MS. Nature Protocols 8, 17–32.

Wei R, Zhang X-C. 2014. Rediscovery of *Cystoathyrium chinense* Ching (Cystopteridaceae): Phylogenetic placement of the critically endangered fern species endemic to China: Phylogenetic placement of *Cystoathyrium chinense*. Journal of Systematics and Evolution 52, 450–457.

Wen B, Mei Z, Zeng C, Liu S. 2017. metaX: a flexible and comprehensive software for processing metabolomics data. BMC Bioinformatics 18, 183.

Wu T, Hu E, Xu S, et al. 2021. clusterProfiler 4.0: A universal enrichment tool for interpreting omics data. The Innovation 2, 100141.

Wu Z-G, Jiang W, Tao Z-M, Pan X-J, Yu W-H, Huang H-L. 2020. Morphological and stage-specific transcriptome analyses reveal distinct regulatory programs underlying yam (Dioscorea alata L.) bulbil growth. Journal of Experimental Botany 71, 1899–1914.

Xu J, Li Q, Yang L, Li X, Wang Z, Zhang Y. 2020. Changes in carbohydrate metabolism and endogenous hormone regulation during bulblet initiation and development in Lycoris radiata. BMC Plant Biology 20, 180.

Yang P, Xu L, Xu H, Tang Y, He G, Cao Y, Feng Y, Yuan S, Ming J. 2017. Histological and Transcriptomic Analysis during Bulbil Formation in Lilium lancifolium. Frontiers in Plant Science 8, 1508.

Young MD, Wakefield MJ, Smyth GK, Oshlack A. 2010. Gene ontology analysis for RNA-seq: accounting for selection bias. Genome Biology 11, R14.

Zhang X. 2012. Lycophytes and Ferns of China. Beijing:Peking University Press.

Zhang Y, Yu BS, Chen Y. 1994. Anatomical Study on the Generation of the Buibil of the Chinese Yam. Journal of China Agricultural University 20, 413–418.

Zhao Y. 2010. Auxin biosynthesis and its role in plant development. Annual review of plant biology 61, 49–64.

Zhao Q. 2016. Lignification: Flexibility, Biosynthesis and Regulation. Trends in Plant Science 21, 713–721.

